# Semi-parametric Empirical Bayes Method for Multiplet Detection in snATAC-seq with Probabilistic Multi-omic Integration

**DOI:** 10.1101/2025.10.23.684138

**Authors:** Yuntian Wu, Haoran Hu, Wei Chen, Johann E. Gudjonsson, Lam C. Tsoi, Xiaoquan Wen

## Abstract

Multiplets, formed when multiple cells are captured in a droplet, produce hybrid molecular profiles that confound single-cell analyses. Detecting multiplets in single-nucleus ATAC-seq (snATAC-seq) data is particularly challenging due to sparsity and overdispersion of chromatin accessibility measurements. We introduce SEBULA, a semi-parametric empirical Bayes model that yields well-calibrated posterior probabilities for multiplet detection, enabling principled false discovery rate control. SEBULA also integrates probabilistic evidence with complementary signals from other modalities, such as scRNA-seq. Benchmarking on simulations and seven annotated trimodal DOGMA-seq datasets demonstrates SEBULA’s superior performance. The open-source software is computationally efficient.

## Background

High-throughput single-cell technologies have transformed molecular profiling by enabling the simultaneous measurement of gene expression, chromatin accessibility, and protein abundance across thousands of individual cells [1]. The emergence of multiomic platforms such as DOGMA-seq [2], CITE-seq [3], TEA-seq [4], SNARE-seq [5], and SHARE-seq [6] has further expanded the field by providing rich, integrated datasets that offer comprehensive views of cellular identity and regulatory mechanisms. However, a persistent technical challenge is the formation of multiplets, where two or more cells or nuclei are co-encapsulated within a single droplet and subsequently processed as one [7], [8], [9]. If not accurately identified and removed, multiplets generate hybrid transcriptomic and epigenomic profiles that can distort downstream analyses, including clustering, differential expression, trajectory inference, and allele-specific accessibility, leading to misleading or spurious results [9], [10], [11].

Multiplets are typically classified as either heterotypic or homotypic, depending on whether the constituent cells originate from distinct or similar cell types or states [12], [13]. Heterotypic multiplets often exhibit intermediate or aberrant expression profiles and are generally more amenable to computational detection. In contrast, homotypic multiplets can be nearly indistinguishable from true singlets, particularly in cell-type-enriched datasets such as those dominated by T cells, where homotypic events may be prevalent yet remain particularly difficult to detect using existing approaches [10], [14].

A variety of computational methods have been developed to identify and remove multiplets, most of which were originally designed for single-modality datasets. For scRNA-seq, tools such as Scrublet [12], scds [15], DoubletFinder [13], solo [16], and scDblFinder [14] simulate artificial doublets *in silico* and train classifiers using low-dimensional embeddings and nearest-neighbor graphs. These methods perform well in transcriptomic data but rely on assumptions about feature structure that do not translate naturally to chromatin accessibility. Moreover, homotypic doublets, formed by cells of the same type or with highly similar transcriptional states, are especially challenging to detect from RNA profiles alone, since they can closely resemble true singlets. [13]. Single-nucleus ATAC-seq (snATAC-seq) data present some unique challenges: they are sparse, capture discrete accessibility states (e.g., 0 for closed chromatin, 1 for open on one parental chromosome, and 2 for open on both), and often exhibit scattered, low-coverage signals. To address these limitations, AMULET [11] introduced a read-level, simulation-free strategy that leverages the genomic loci with high-coverage to detect both heterotypic and homotypic multiplets. While interpretable and efficient, AMULET assumes a homogeneous singlet population and imposes strong distributional assumptions, which may restrict its performance in datasets with heterogeneous sequencing depth, variable sample composition, or intrinsic overdispersion.

Complementary experimental and genotype-based approaches, such as Cell Hashing [17], MULTI-seq [18], and demuxlet [19], can identify inter-sample multiplets and provide orthogonal bench-marks. However, these methods require additional experimental steps and costs, cannot be applied retrospectively to unlabeled datasets, and often fail to detect within-sample (same-donor) homotypic events. As a result, computational strategies that operate directly on single-cell data, especially snATAC-seq data, remain indispensable for comprehensive multiplet detection.

The advent of multiomic assays has enabled the simultaneous profiling of transcriptomic and epigenomic features within the same cell, creating new opportunities for enhanced multiplet detection through signal integration. COMPOSITE [20] was recently introduced as a unified framework that models stable features from RNA, ATAC, and protein modalities within a likelihood-based framework, demonstrating enhanced performance by leveraging complementary information. Nevertheless, alternative inference strategies that integrate evidence across data modalities, particularly those allowing direct incorporation of established, modality-specific detection tools, remain valuable. Such flexible computational approaches not only accommodate diverse data types but also support continued progress in modality-specific detection methods through a divide-and-conquer paradigm.

Here, we present SEBULA, a novel computational method for multiplet detection in single-nucleus ATAC-seq data. SEBULA is built on a semi-parametric inference framework to achieve improved sensitivity and specificity over existing approaches. To extend this applicability, we introduce a general probabilistic strategy that integrates evidence across diverse features and modalities, enabling robust multiplet detection in multiomic single-cell datasets. SEBULA is released as an open-source software package and is freely available at https://github.com/Yuntian0716/SEBULA.

## Results

### Overview

SEBULA implements a semi-parametric probabilistic model for multiplet detection in snATAC-seq data (Figure 1). Following a preprocessing strategy similar to AMULET, we generate a binary matrix of multi-read sites, where rows corresponds to a genomic loci with more than two overlapping reads in at least one cell (i.e., a high-coverage locus), and columns represent individual nuclei. From this matrix, we compute the high-coverage locus count (HCLC) for each cell, which serves as input to SEBULA. Intuitively, cells with unusually high HCLC values are more likely to be multiplets.

**Figure 1.**
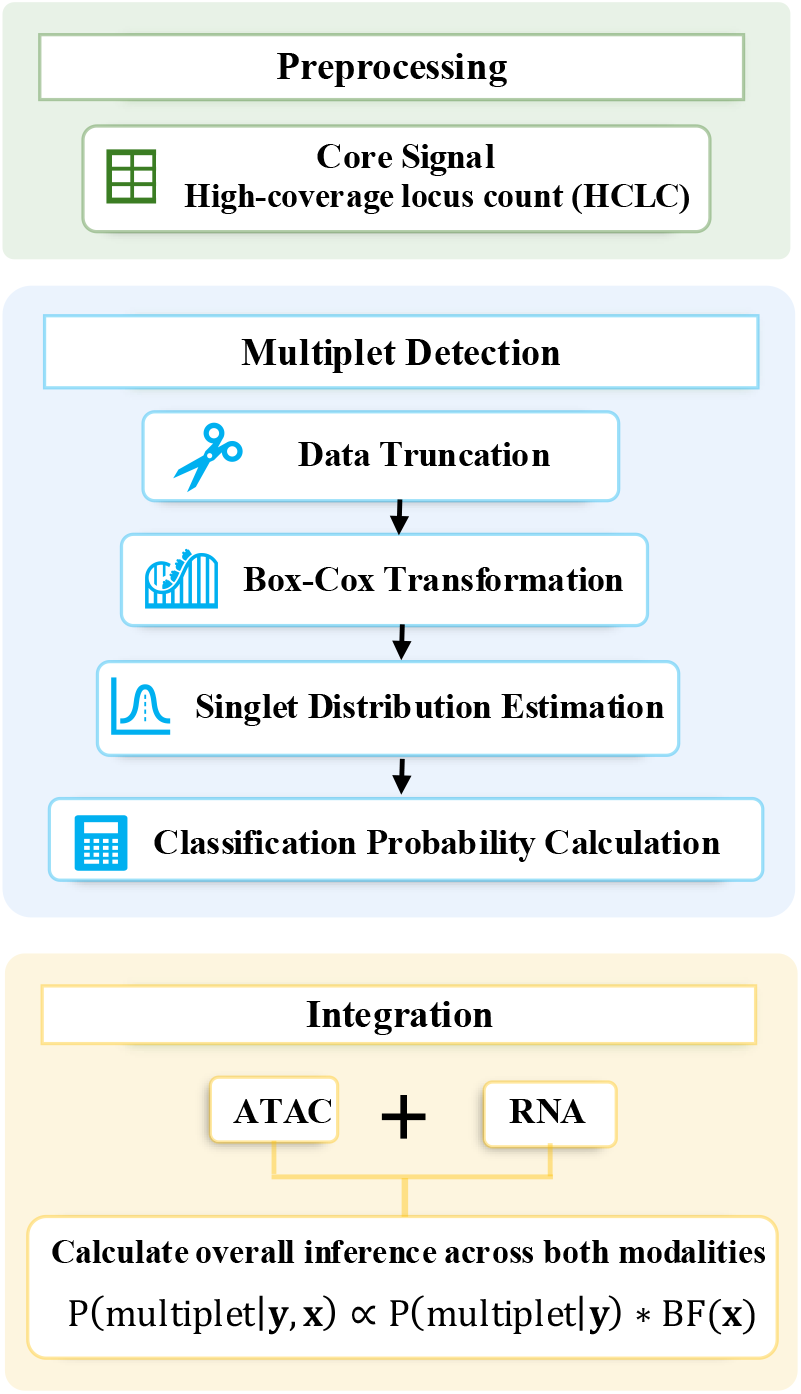
Overview of the SEBULA framework for multiplet detection and multisource integration. The SEBULA workflow consists of three modular stages: (i) preprocessing, (ii) chromatin-based multiplet detection, and (iii) multi-source integration. *Top panel (Preprocessing):* For each cell, we compute the high-coverage locus count (HCLC), defined as the number of genomic regions with *>*2 overlapping ATAC-seq reads. This biologically motivated metric captures local read enrichment, which is elevated in multiplets due to the presence of multiple nuclei. *Middle panel (Multiplet Detection):* Raw HCLC distributions are truncated to remove low-information cells and then normalized using a Box–Cox transformation to reduce skewness. The singlet distribution is then estimated via a semi-parametric central matching strategy applied to the central region of the transformed data, where singlets are expected to dominate. Each cell receives a posterior probability of being a singlet, with multiplets identified as low-probability outliers. *Bottom panel (Integration):* SEBULA supports flexible incorporation of complementary data, such as classification probabilities from RNA-based methods (e.g., scDblFinder). These signals are combined using a Bayesian framework, enabling across modalities or pipelines to improve multiplet classifications.

The key innovation of SEBULA lies in its semi-parametric procedure for estimating the distribution of HCLC in singlet nuclei directly from observed data. By contrast, methods such as AMULET, assume parametric models, for example, that the number of multi-read sites in singlets follows a Poisson distribution. However, analyses of snATAC-seq datasets with groundtruth multiplet annotations show that commonly used count models, such as the Poisson and negative binomial distributions, fit poorly to empirical data, resulting in suboptimal performance in multiplet detection (Figure 2).

**Figure 2.**
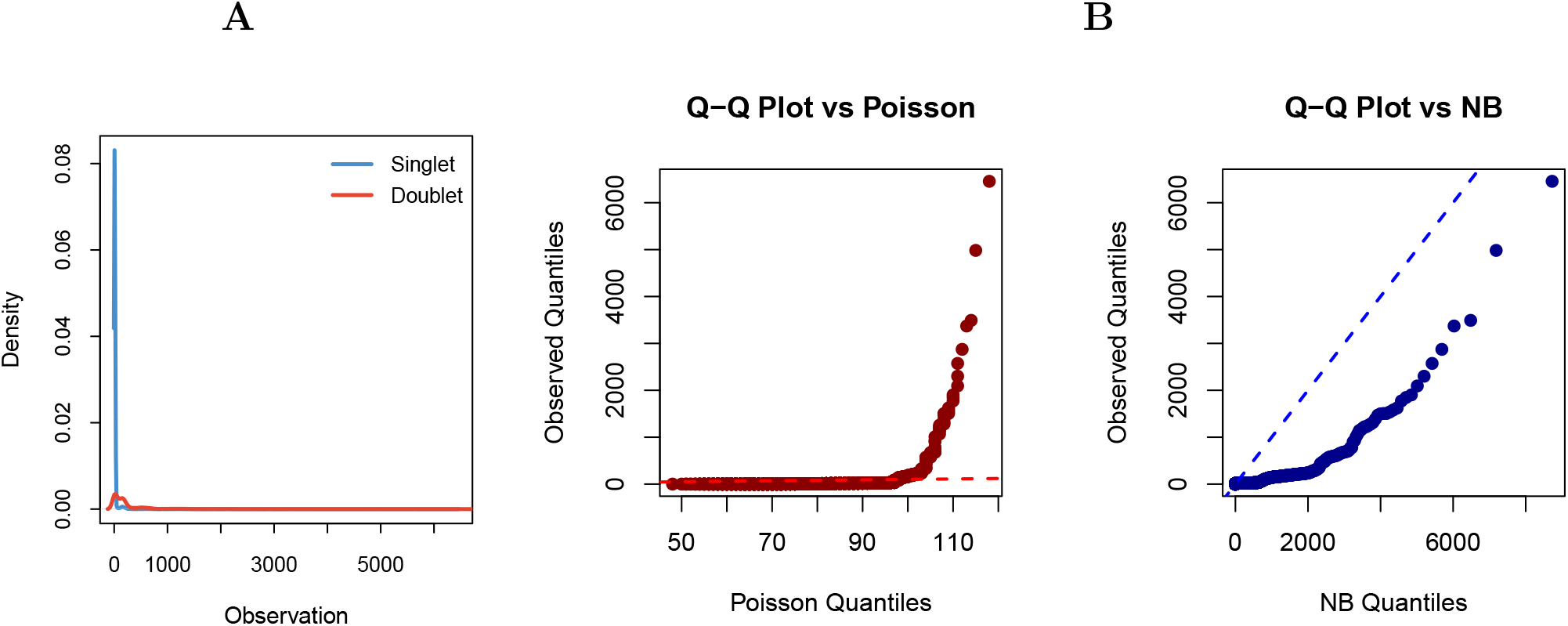
Deviations from Poisson and negative binomial assumptions in highcoverage locus counts. **A** Density plot of the number of genomic loci with *>*2 overlapping fragments per nucleus in a DOGMA-seq dataset (PB-3), stratified by ground truth singlet and doublet labels obtained from cell hashing. The singlet distribution (light blue) is heavily right skewed and multimodal, suggesting deviation from standard parametric assumptions. **B** Quantile–quantile (Q–Q) plots comparing the observed singlet distribution (after excluding known doublets) with theoretical quantiles under Poisson (middle) and negative binomial (right) models. Model parameters were estimated by the method of moments across the full cell population. The Poisson model underestimates tail density due to it equidispersion assumption, while the negative binomial model better captures dispersion but fails to model extreme tails and multimodality, highlighting limitations of both distributions in modeling singlet variation. These limitations motivate a semi-parametric approach to singlet modeling that relaxes rigid distributional assumptions.

Informed by these observations, we reason that effective multiplet detection should prioritize distinguishing multiplets from singlets with moderate to high HCLC, since cells with extremely low counts are almost certainly singlets and thus less informative for multiplet detection. To achieve this, we adopt a strategy analogous to the *zero assumption* in non-parametric empirical Bayes (NPEB) inference [21], enabling accurate characterization of the right-tail behavior of the singlet distribution. Specifically, cells with very low HCLC (threshold user-defined) are excluded, and the remaining counts are transformed into a continuous domain using a rank-preserving Box-Cox transformation. The empirical density of the transformed values, denoted by *f* (*x*), it then estimated and modeled as a mixture of singlet and multiplet distributions:

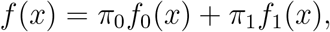

where *x* denotes the transformed count, and *π*_0_ and *π*_1_ are the proportions of singlets and multiplets, respectively.

Following the principle of the zero assumption, we make two key assumptions: (i) singlets are substantially more abundant than multiplets (*π*_0_ *≫ π*_1_), and (ii) cells near the mode of the empirical distribution are predominantly singlets. Leveraging these assumptions, we apply non-parametric density estimation around the mode of *f* (*x*) to infer the singlet distribution *f*_0_(*x*) and estimate its mixing proportion *π*_0_ (Methods). The multiplet distribution *f*_1_(*x*) and its mixing proportion *π*_1_ are subsequently induced. With these distributional estimates, the posterior probability that a cell is a multiplet is computed via Bayes’ rule:

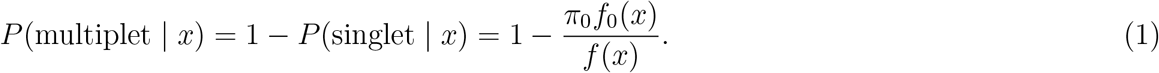

These estimates also enable a principled framework for local and Bayesian FDR control procedure in multiplet detection. Specifically, under a decision rule that classifies all cells with HCLC ≥ *t* as multiplets, the corresponding estimated false discovery rate (FDR) is given by

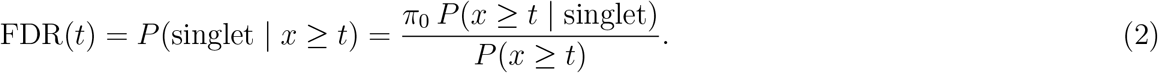

Quantifying multiplet evidence from the HCLC on a probabilistic scale allows seamless integration with other probabilistic signals, whether derived from additional features in snATAC-seq data, or from complementary modalities such as scRNA-seq. Consequently, SEBULA can be combined with other multiplet detection tools that generate probabilistic outputs, including scDblFinder and COMPOSITE, to further enhance sensitivity and specificity.

This integration follows the principle of Bayesian sequential updating [22]. Specifically, the Bayes factor for the HCLC data, BF(***x***), can be obtained directly from the posterior probability (1) given the snATAC-seq HCLC data ***x***. By treating the posterior probability derived from additional feature data **y**, *P* (multiplet | **y**), as a feature-informed prior, we construct a naïve Bayes classifier that integrates evidence across both sources. This yields the updated classification posterior:

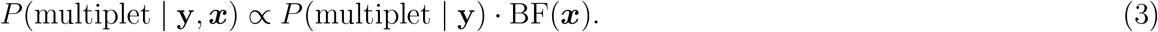

A detailed derivation is provided in the Methods section.

### Evaluation using Simulated Artificial Doublets

We generated synthetic snATAC-seq data to benchmark the performance of SEBULA and AMULET using only HCLC-derived information. Starting from a real DOGMA-seq dataset from Xu *et al*. [2], we extracted the scATAC-seq modality and first removed all annotated multiplets. Artificial doublets were then created by randomly pairing the remaining 15,788 singlet cells, thereby preserving the authentic propoeties of true singlets. To assess performance across various settings, we constructed multiple datasets with artificial doublets proportions from 5% to 25%. For each simulated dataset, the resulting multi-read site matrix and HCLC information was provided as input to both SEBULA and AMULET.

We evaluated the sensitivity and specificity of each method using receiver operating characteristic (ROC) curves and F1 scores across all simulated datasets. SEBULA consistently outperformed AMULET by a wide margin under all conditions (Figure 3 and Supplementary Figure S1). Notably, AMULET’s performance declined as the proportion of doublets increased, likely due to its reliance on a global mean to approximate the singlet distribution. This approach makes AMULET sensitive to distortion when doublets are abundant, as their inclusion inflates the global average and miscalibrates the null model. In contrast, SEBULA remained robust across varying doublet frequencies, maintaining strong discrimination and better calibration, as reflected by consistently higher area under the ROC curve (AUC) (Supplementary Figure S1).

**Figure 3.**
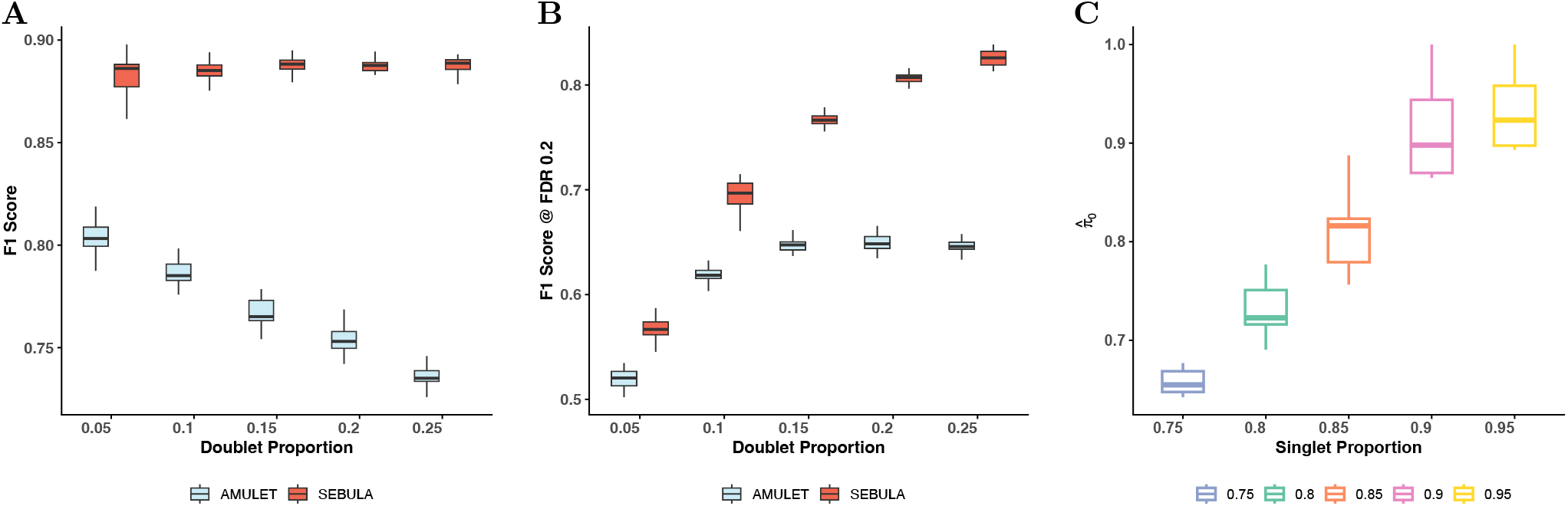
Performance evaluation of SEBULA and AMULET across simulated doublet scenarios. **A** Optimal F1 score achieved across a range of thresholds for each simulated doublet proportion (5% to 25%). **B** F1 score evaluated at a fixed decision threshold of FDR *<* 0.2 (right-sided Bayesian FDR for SEBULA and adjusted *p*-value for AMULET). **C** Accuracy of singlet proportion estimates 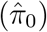 across varing true singlet proportions. In all panels, each boxplot summarizes results from 20 replicate simulations per condition.

To better approximate real-world application, we allowed both methods to determine classification cutoffs by estimating the false discovery rate (FDR) at the 20% level, with AMULET using built-in *q*-values and SEBULA applying Bayesian FDR control (Eqn (2)). As shown in Figure 3B, SEBULA continues to consistently outperform AMULET in terms of F1 scores. Furthermore, SEBULA’s estimates of the multiplet prevalence were reasonably accurate under our simulation settings (Figure 3C), suggesting that these estimates could serve as a useful quantitative indicator of snATAC-seq data quality.

We also implemented and evaluated an alternative parametric mixture model in which the HCLC distribution in singlets is modeled with a negative binomial distribution. We refer to this approach as the negative binomial mixture model (NBMM), which can be considered as an intermediate approach between AMULET and SEBULA (Supplementary Material). However, when applied to the simulated artificial doublet data, NBMM exhibited poor calibration of posterior classification probabilities: both the estimated singlet proportions and the posterior probabilities for the vast majority of cells are heavily concentrated near 1 across all simulation settings (Supplementary Figure S2). We interpret this as evidence that the negative binomial assumption is inadequate for modeling singlet distributions in real data. While the negative binomial distribution addresses some limitations of the Poisson model used by AMULET (such as the inability to handle overdispersion or zero-inflated counts), it appears overly permissive for larger HCLC values and thus fails to effectively separate multiplets from singlets. Viewed from another perspective, this underscores SEBULA’s key advantage: it’s semi-parametric strategy provides a more flexible and accurate model of singlet distribution in real data.

### Multiplet Detection in Trimodal Single-Cell Data

We applyed SEBULA to seven trimodal single-cell datasets generated by DOGMA-seq experiments and benchmark its performance against state-of-the-art multiplet detection methods. DOGMA-seq enables simultaneous profiles the transcriptome (scRNA-seq), chromatin accessibility (scATAC-seq), and cell surface protein expression (via antibody-derived tags, or ADTs) in single cells. While SEBULA directly leverages only the scATAC-seq modality, these multimodal datasets enable evaluation of ensemble strategies that integrate SEBULA’s ouput with classification probabilities derived from additional modalities. The seven benchmarking datasets were all generated from experiments on human peripheral blood T cells, including one dataset from Xu *et al*. [2] (GSE200417) and six from Hu *et al*. [20]. In all cases, high-confidence binary labels for multiplet classification were obtained from cell hashing and are treated as ground truth for evaluation.

Among the methods that reply solely on the scATAC-seq modality, SEBULA consistently outperforms AMULET and ArchR in terms of F1 scores and performs comparably to scDblFinder (Figure 4). Notably, both ArchR and scDblFinder classify multiplets using in silico doublet generation and leverage multiple features from scATAC-seq data, excluding HCLC. The performance gap between SEBULA and AMULET corroborates the findings from our synthetic data simulations. We further hypothesize that the benchmarking datasets are enriched for homotypic multiplets, a setting in which the HCLC statistic are particularly effectivey, likely explaining SEBULA’s advantage over ArchR and, in some cases, over scDblFinder.

**Figure 4.**
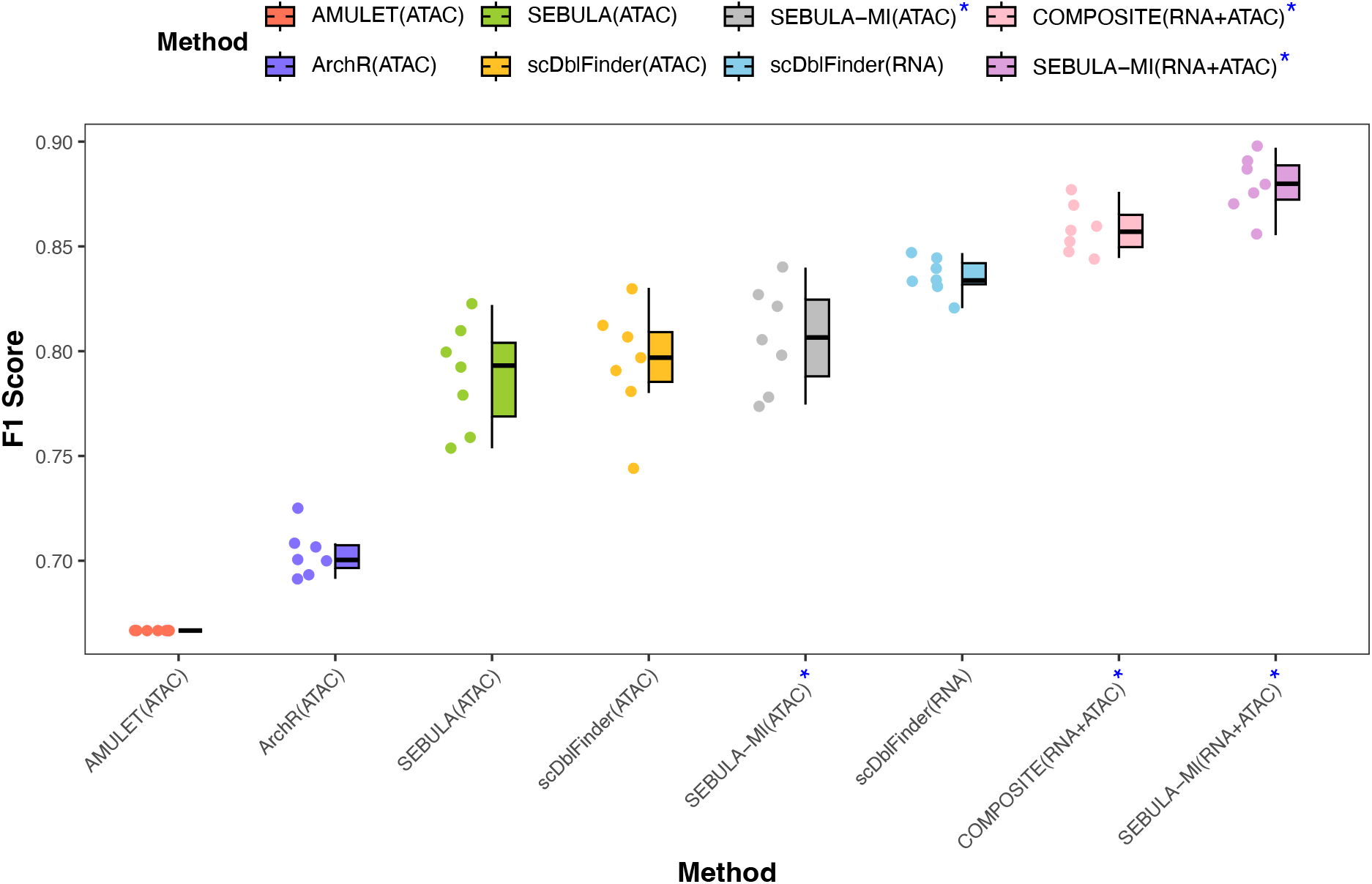
Benchmarking SEBULA and SEBULA-MI against existing multiplet detection methods using ground-truth annotations. Boxplots showing F1 score performance of each multiplet detection method evaluated across seven DOGMA-seq datasets with experimentally validated multiplet labels derived from cell hashing. Methods include AMULET, ArchR, and scDblFinder (applied to RNA and ATAC modalities), the COMPOSITE multiomic framework, and our proposed method, SEBULA, along with its multi-source integration extension, SEBULA-MI. SEBULA (green) and SEBULA-MI (gray for ATAC-only) consistently outperformed AMULET (red) and ArchR (blue), and achieved comparable or superior accuracy to scDblFinder (yellow for ATAC-only). Notably, SEBULA-MI (purple) achieved the highest overall performance across all datasets. Methods denoted with a blue asterisk (*) incorporate multiple sources of information (e.g., RNA+ATAC). Each point represents a sample-level evaluation; boxes span the interquartile range with the median marked by a horizontal line, and whiskers extend to 1.5× the IQR.

To test this, we applied the scheme outlined in Eqn (3) to integrate the classification probabilities from scDblFinder ATAC-seq analysis with those from SEBULA. The ensemble effectively combines the features leveraged by scDblFinder with the HCLC statistics captured by SEBULA. The ensemble consistently improved upon both individual methods, achieving the best overall performance within the ATAC-seq modality across all benchmarking datasets (Figure 4).

We next extended this strategy across modalities by combining scRNA-seq based inference from scDblFinder with ATAC-seq–based inference from SEBULA, again using Eqn (3) to compute per-cell classification probabilities. Notably, scRNA-seq–based inference alone achieved higher sensitivity and specificity for multiplet detection than scATAC-seq–based inference. Nevertheless, integrating evidence from both modalities yielded clear improvements over using either modality alone, underscoring the complementary value of multimodal data. Remarkably, the red performance of the ensemble classifier is even on par (or slightly better than) COMPOSITE, a state-of-the-art method specifically designed for cross-modality multiplet detection.

## Discussion

SEBULA is designed for multiplets detection in snATAC-seq data. Like AMULET, it focuses on a single feature, HCLC, which we demonstrate to be highly informative for distinguishing multiplets from singlets. The main contribution of this work is the semi-parametric modeling of the singlet distribution within an empirical Bayes framework. Because of the complexity of scATAC-seq experiments and the construction of the HCLC statistic (which involves explicit censoring of genomic loci), standard parametric statistical models are difficult to fit adequately to singlet data. Consequently, nonparametric strategie such as artificial doublets simulation have gained popularity. However, methods based on synthetic doublets (e.g., scDblFinder and ArchR) may suffer from reduced sensitivity when the true multiplet proportion is high. In contrast, SEBULA estimates the singlet distribution by matching the mode of the empirical distribution of Box-Cox–transformed HCLC values. This strategy, motivated by the central limit theorem under the assumption that singlets dominate the population, is shown to be robust and effective across a wide range of multiplet proportions in both our synthetic and real datasets.

By leveraging the HCLC statistic, SEBULA detects homotypic multiplets as effectively as heterotypic ones, addressing a key performance gap in practical multiplet removal. This point is also discussed in [11]. More importantly, we propose a computational framework for integrating SEBULA’s probabilistic evidence of multiplet with outputs from other detection tools. In our DOGMA-seq data analysis, we find that these different sources of evidence are individually complementary and collectively more powerful, underscoring SEBULA’s effectiveness in handling complex multimodal data beyond single-modality scATAC-seq. Our feature integration strategy exemplifies a general divide-and-conquer approach to accommodating ever increasingly complex genomic data structure: rather than attempting to jointly model all features across modalities simultaneously, it can be more efficient to design targeted analysis modules that maximize the utility of individual features, and then probabilistically combine their evidence. Nonetheless, we acknowledge that integration across multiple features and modalities remain an open challenge, and we view SEBULA as a foundation on which more sophisticated multimodel integration strategies can be built in future work.

Following standard practice in the field, our performance evaluation of SEBULA primarily emphasizes the *ranking* of likely multiplets based on classification probabilities, with sensitivity and specificity assessed across a range of thresholds. However, we argue that equal or greater attention should be paid to the calibration of the classification probabilities themselves. While accurate estimation of the overall multiplet proportion, as shown in our synthetic data experiments, provides indirect evidence of proper calibration, direct evaluation is necessary both for a more complete assessment and for offering practical guidance in selecting decision thresholds. We also advocate distinguishing this form of evaluation from formal hypothesis testing, where singlets are treated as the null class. Although the two frameworks may appear similar, the consequences of misclassification differ: in downstream analysis, misclassifying a singlet as a multiplet (false positive) arguably is not inherently more detrimental than misclassifying a multiplet as a singlet (false negative).

## Conclusions

SEBULA provieds an efficient and robust computational framework for detecting and annotating multiplets using only read count information from scATAC-seq data. Across extensive simulations and real datasets, it consistently demonstrates improved sensitivity and specificity with existing methods. Moreover, SEBULA’s probabilistic output can be readily integrated with complementary evidence from other diverse features and modalities, enabling even more effective multiplet detection in multimodal settings. The SEBULA software is open-source and freely available at https://github.com/Yuntian0716/SEBULA.

## Methods

### Pre-processing and HCLC Generation

To generate the high-coverage locus count (HCLC) statistics required for SEBULA, we largely follow the preprocessing procedures established in AMULET, which consist of two main stages.

In the first stage, we process fragment files containing all read alignments along with a set of confidently called cell barcodes. For each autosomal chromosome in each cell, we retain fragments with insert sizes less than 900 base pairs. We then identify and record high-coverage loci – defined as genomic intervals overlapped by more than two fragments within a single cell – using a sweepline algorithm.

In the second stage, we aggregate the high-coverage loci identified across all cells to create a non-redundant set of genomic intervals for each chromosome. This is achieved using a greedy interval-merging algorithm. Based on these merged loci, we construct a binary matrix in which each row corresponds to a high-coverage locus and each column corresponds to a cell. An entry in the matrix is set to 1 if the cell contains a fragment overlapping the corresponding locus, and 0 otherwise. The HCLC for each cell is then computed as the sum of entries in the corresponding column.

### Semi-parametric Modeling of HCLC

Let ***X*** = *x*_1_, …, *x*_*N*_ denote the HCLC values for *N* cell barcodes in an experiment. We first exclude cells with HCLC strictly less than a predefined threshold *x*_min_ under the assumption that such cells are predominantly singlets. In the Method part, we provide a computational procedure for selecting an optimal value of *x*_min_ based on the overall goodness-of-fit of the SEBULA model. However, we find that SEBULA’s results are generally robust when *x*_min_ ∈ [0, *Q*_1_], where *Q*_1_ denotes the 25th percentile. We denote the set of filtered HCLC values by ***X***_*T*_ and the number of retained cells by *N*_*T*_. For each *x*_*j*_ ∈ ***X***_*T*_, we apply a variance-stabilizing and rank-preserving Box–Cox transformation [23],

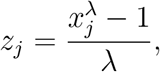

where the parameter *λ* is determined by maximizing the Box–Cox log-likelihood. SEBULA models the Box–Cox transformed HCLC data, ***Z***_*T*_, using a two-component mixture distribution:

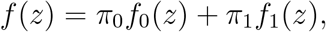

where *f*_0_ is assumed to be a normal density with (unknown) mean, *δ*_0_, and variance, 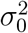.

### Mixture Density Estimation via Poisson Spline Smoothing

We estimate the mixture density *f* (*z*) using a smoothed histogram of the truncated variable ***Z***_*T*_, partitioned into *K* equally spaced bins (default *K* = 120). For *i* = 1, …, *K*, let *y*_*i*_ denote the observed count in the *i*th bin and *c*_*i*_ the center of the bin.

Following Efron [21], [24], [25], we treat *y*_*i*_ as independent Poisson observations and fit a generalized linear model (GLM) with natural spline basis of degree 7. Specifically, we assume

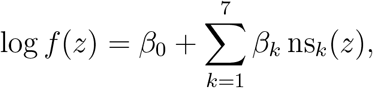

where ns_*k*_(*z*) denotes the *k*th basis function of the natural spline. This yields a smooth estimate 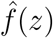 of the mixture density. The fitted values at bin centers, 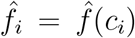, are normalized to produce a discrete approximation of the density:

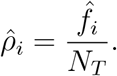

A smooth estimate of the cumulative distribution function (CDF), 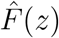, is subsequently obtained by linearly interpolating the bin centers *{c*_*i*_*}* against the cumulative sums of 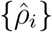.

### Singlet Density Estimation by Central Matching

We apply Efron’s central matching method [21] to estimate the singlet density *f*_0_(*z*) and the singlet proportion *π*_0_ in the transformed HCLC data. The key assumption here is that the vast majority of the transformed HCLC values within a predefined central window correspond to singlet cells. (By default, we specify the central window spanning the 20th to 60th percentiles of the empirical distribution of ***Z***_*T*_.) By isolating data within this range, we fit a quadratic model to the log of the sub-density 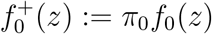 as follows:

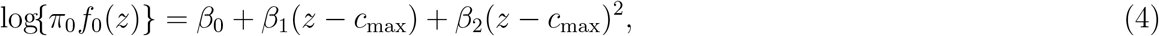

where *c*_max_ is the center of the histogram bin where the estimated mixture density 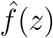 attains its maximum.

Assuming 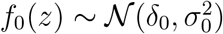, the log sub-density expands as

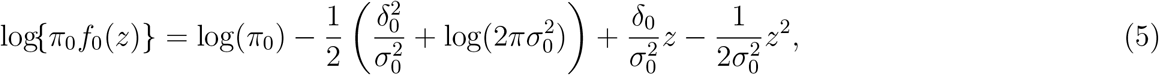

The least squares estimates, 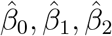, from the fit of (4) induce the estimates of 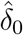 and 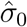 for the normal density of the singlet distribution:

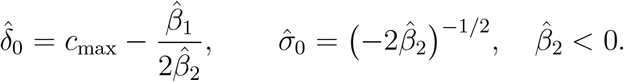

An unnormalized estimate of the singlet count in each histogram bin *i* is then computed by

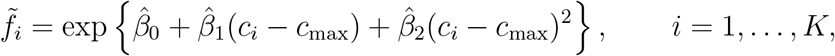

where 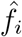 denotes the fitted count from the Poisson spline model at the *i*th bin center, *c*_*i*_. Finally, the singlet proportion *π*_0_ is estimated by normalizing the total fitted singlet density:

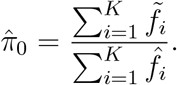

Note that 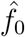 represents the proportion of singlets among the cells after initial filtering. To report the estimate of the singlet proportion in the *entire* original library, we use

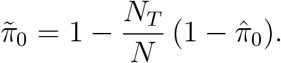

### Optimizing Truncation Point *x*_min_

In practice, we find that filtering out zero and low values of HCLC can substantially improve the efficiency of mixture distribution estimation. Although the final results for multiplet classification are generally robust to the specific choice of the threshold *x*_min_, provided that *x*_min_ ≥ 0, we propose a simple computational procedure to aid in selecting an *x*_min_ value to optimize the performance of SEBULA.

The proposed procedure evaluates the goodness-of-fit between the transformed HCLC values within a specific range and the estimated normal distribution using cross-entropy loss. Specifically, after filtering the data and fitting the mixture model, we define

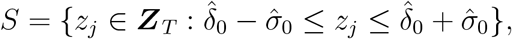

and compute the cross-entropy for the corresponding *x*_min_ as

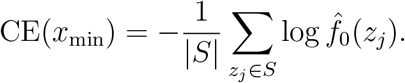

The cross-entropy measures the discrepancy between the fitted normal distribution and the empirical distribution of the data; lower values indicate a better fit. Therefore, selecting an *x*_min_ that minimizes the cross-entropy supports the validity of the central matching assumption.

Because the mixture fitting procedure is computationally efficient, SEBULA can quickly evaluate cross-entropy values over a range of *x*_min_ settings. To balance improved goodness-of-fit with retaining sufficient data for stable estimation of the mixture model, SEBULA constructs a trace plot of CE(*x*_min_) and selects the elbow point as the optimal threshold (Supplementary Figure S3).

### Multiplet Classification

Based on the overall procedure, we compute the posterior probability that cell *j* is a multiplet as

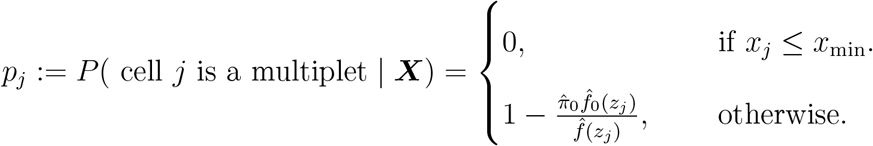

Thresholding the *p*_*j*_ values is a standard classification approach in machine learning practice. By default, we classify cells with *p*_*j*_ *>* 0.8 as multiplets, which corresponds to applying a local false discovery rate threshold of 20%.

Alternatively, we also implement a classification procedure that controls the overall false discovery rate (FDR), primarily for comparison with AMULET. Note that the estimated densities 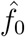 and 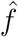 induce corresponding cumulative distribution functions 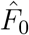 and 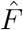. For a decision rule that classifies all cells with Box–Cox transformed values *z* ≥ *t* as multiplets, the estimated FDR at threshold *t* is given by

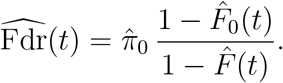

This approach allows users to pre-specify a target FDR level (e.g., 20%) and then identify the corresponding cutoff value of *z* directly from the data.

### Probabilistic Evidence Integration

To integrate probabilistic evidence from HCLC with evidence derived from other data features and/or modalities, we adopt a general Bayesian framework for sequentially updating posterior multiplet probabilities.

We regard the existing posterior probability for a target cell, *P* (multiplet | ***y***), as the new prior for re-analyzing the HCLC data. We then recover the HCLC likelihood for each cell from the earlier HCLC-only analysis. Specifically, we compute the marginal likelihood of the HCLC data, i.e., the Bayes factor, for cell *j* as

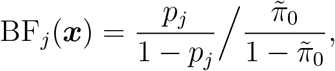

where *p*_*j*_ is the posterior multiplet probability based on HCLC alone, and 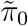 is the estimated proportion of singlets in the original library.

Assuming conditional independence between ***x*** and ***y***, i.e.,

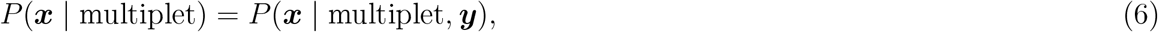

the updated posterior probability incorporating HCLC evidence is given by the Bayes rule:

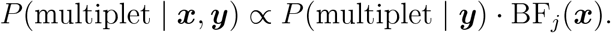

Note that the conditional independence assumption in (6) is equivalent to the naive independence assumption used in the naive Bayes classifier. While this assumption may not hold exactly in practice, empirical evidence suggests that it is approximately valid, particularly when integrating data across distinct modalities.

## Declarations

### Ethics approval and consent to participate

Not applicable.

### Consent for publication

Not applicable.

### Availability of data and materials

All datasets analyzed in this study were previously published and are publicly available:

The DOGMA-seq dataset is available from the Gene Expression Omnibus (GEO) under accession number GSE200417. The six PBMC datasets (“PB-1”, “PB-2”, “PB-3”, “PB-4”, “PB-8”, “PB-9”) were generated and published in the COMPOSITE study (Hu et al., 2024), and are publicly available at: https://doi.org/10.5281/zenodo.11167173.

To run SEBULA, we additionally used filtered and raw feature-barcode matrices (filtered_feature _bc_matrix.h5, raw_feature_bc_matrix.h5) and ATAC fragment files (atac_fragments.tsv.gz), which were prepared and shared directly by the dataset author Haoran Hu. The SEBULA software is freely available at: https://github.com/Yuntian0716/SEBULA

### Authors’ contributions

YW developed the SEBULA model, implemented the software, performed benchmarking analyses, and drafted the manuscript. HH prepared and shared the COMPOSITE benchmarking datasets, and along with WC, provided constructive feedback on the methodology and results. JEG contributed biological interpretation and provided critical feedback on study design and manuscript revisions. LCT and XW supervised the project from biological and statistical perspectives, respectively. All authors contributed to manuscript revision and editing. All authors read and approved the final manuscript.

